# DEGRADATIVE POTENTIALS OF BACTERIA ISOLATED FROM CRUDE OIL CONTAMINATED SOIL

**DOI:** 10.1101/2023.10.16.562498

**Authors:** Jennifer Ada Augustine

## Abstract

Petroleum and its products continue to serve as a principle source of energy for industries and daily life. However, their release into the environment is a worldwide concern since some products are acutely toxic or possess mutagenic, teratogenic and carcinogenic properties. Several oil disposal methods have been applied over time with remediation emerging as the most promising technology. It takes advantage of the versatility of soil microbes to degrade hydrocarbon contaminants. Unlike conventional disposal methods, bioremediation ids an environmentally friendly and cost effective method that stimulate natural processes for complete degradation of hydrocarbons into innocuous compounds. This study focused on isolation, morphological and biochemical characterization of bacteria possessing hydrocarbon degrading properties. The study also aimed at optimizing appropriate culture conditions for the isolates. Isolation of hydrocarbon degrading bacteria from soils polluted with crude oil in Elebel, Emeyal I, Emeyal II in Ogbia, Bayelsa State was carried out using mineral salt medium supplemented with 1% crude oil, and the isolates were screened on their ability to use carbon as their sole source of carbon or alternative pathway. Characterization of the isolates was carried out by performing Gram’s iodine test, catalase, oxidase, indole, citrate, motility, coagulase tests. The degrading potential of the isolates were obtained by subculturing them in mineral salt medium for two weeks at room temperature, organisms that could utilize all crude oil made available to them were considered strong degraders while those that could not utilize all the crude oil made available to them were considered weak degraders. Among 40 microbes isolated, 30 were considered strong degraders while 10 were considered weak degraders and these isolates included *Pseudomonas* spp, *Enterobacter* spp, *Escherichia coli*, *Arthrobacter* spp, *Bacillus* spp, *Acinetobacter* spp, *Staphylococcus aureus*, coagulase negative *Staphylococcus* spp, *Klebsiella* spp, *Micrococcus* spp, *Vibrio* spp. The characterized isolates may constitute potential candidates for biotechnological application in environmental cleanup of petroleum contaminants.

## INTRODUCTION

Petroleum oil is a great source of energy that has been known throughout historical time. It is oily, flammable and present in suitable rock strata. Petroleum is a complex mixture of gaseous, liquid and solid hydrocarbons. It occurs naturally beneath the earth’s surface. Petroleum’s hydrocarbons are mainly made of traces of various nitrogenous and sulphurous compounds. It can be extracted and refined to produce fuels including petrol, paraffin, kerosene, gasoline and diesel (USEPA, 2003; Akpor *et al*., 2007).

Petroleum-based products are a principle source of energy for industries and daily life, making them a vital commodity central to the global economy (Jahangeer and Kumar, 2013). These products include petrol. Gasoline, kerosene, diesel oil, lubricating oil amongst others. They originate from crude oil whose main constituents are hydrocarbon compounds derived from ancient algae and plant remains found in reservoirs under the earth’s surface. Petroleum products are divided into four classes: saturates, aromatics, resins and asphaltenes (Tebyanian *et al*., 2013)

All petroleum products are originated from crude oil whose major constituents are hydrocarbons. Crude oil is composed of complex mixtures of paraffinic, alicyclic and aromatic hydrocarbons (Forkan *et al*., 2010). Benzene, Toluene and Xylene (BTX) are major aromatic hydrocarbon in many petroleum products (Anitha, 2009), which contaminated environment was hazardous to human and animals and decreases the agricultural productivity of the soil (Bijay, 2012). Prolonged exposure of BTX may cause the lung, heart, liver and kidney disease, bone marrow damage and benzene has been cause cancer (Mandri and Lin, 2007). These illnesses were affected by direct contact with the contaminated soil, vapors from the contaminants, and from secondary contamination of water supplies within and underlying the soil. Multiple techniques have been developed to resolve the problem of petroleum pollution. Physical and chemical method to reduce hydrocarbon pollution is expensive than biological methods (Esin *et al*., 2011). Among these procedures bioremediation is currently gaining importance. The biodegradation of crude oil by microorganisms is one of the primary ways to remove crude oil from contaminated area by use of microorganisms or by their enzymes (Bhasheer *et al.,* 2014). Some types of bacterial isolates are able to degrade the hydrocarbons and use of them as a source of carbon and energy (Venosa and Xueqing, 2003). Among the different types of hydrocarbons, benzene is of major concern as it is a stable, water-miscible, highly mobile, poisonous, and cancer-causing aromatic compound. Successful degradation of benzene by microorganisms in an aerobic environment has been reported; however, under anaerobic conditions its rate of biodegradation is observed to be very slow and poor, but same time successful in aerobic condition (Chaillan *et al*., 2006).

Petroleum was found in oil pits, sands, seeps, springs, and tar pits. Its early uses were for street paving, medicinal use, lighting, water proofing, mortar as well as fire weapon in defensive warfare (Singh and Lin, 2008). 90% of world oil production was from Azerbaijani region between 1850s-1860s. From that time, investors became interested in the possibility of selling oil commercially. The increased interest led to the discovery of an unremarkable hole, producing about 10 barrels per day. That finding made Pennsylvania ‘oil rush’ responsible for half of world’s oil production until 1901 (O’Neil, 2014).

Hydrocarbon pollution is widely recognized as a serious environmental problem since it’s not only serious damage to living things, it also causes adverse effect on the natural environment and ecosystem. Release of hydrocarbons into the environment, whether accidentally or due to human activities is a main cause of water and soil pollution. Hydrocarbon components have been known to belong to the family of carcinogens and neurotoxic organic pollutants (Nilanjana and Preethy, 2011).

Decades after the discovery of petroleum, petroleum’s use was limited to lighting and lubricating. By the year 1910, people began to realize the true potential of oil. Consequently, fields were developed in Iran, Sumatra, Venezuela, Peru, and Mexico to exploit it. Today, the world is heavily dependent on petroleum oil for motive power, lubrication, fuel, dyes, drugs and many other synthetics. Globalization and the need for transporting people and goods from one place to another increased the use and demand for petroleum. Large amounts of petroleum products needed to be produced every year to continue to supply energy and meet the need of the fast growing society (Hussein *et al.,* 2012).

The increased worldwide demand for petroleum had increased from 63 million barrels per day in 1980 to 85 million barrels per day in 2006. Another increase is projected to increase 37% by 2030 (118 million barrels per day) from 2006 quantity. The direct consequence of that increase is spillage and leakage in the environment. Petroleum can be accidentally or deliberately released into the environment. Approximately 600,000 tonnes metric in 2010 and 4000 tonnes in 2014 of petroleum and petroleum products were lost during exploration, production, refining, transport, distribution, and storage. The wrong channeling and discharge of used oil from operation, contribute immensely to land pollution. Pollution of lands has immensely created serious environmental problems. That often results in huge disorder of both the biotic and abiotic components of the ecosystems (Das and Chandran, 2011; Eze *et al.,* 2014).

Crude oil spill, no matter its quantity and size (minor, medium, major or disaster may cause minor or severe damages to the environment and all forms of life dependent on the environment. The release of crude oil into the environment causes enormous damages to the environment due to the presence of many toxic compounds such as polycyclic aromatic hydrocarbons, benzene and its substituted and cycloalkane rings in relatively high concentration (Agarry and Ogunleye, 2012). Oil spill on land may lead to retardation of vegetation growth and cause soil infertility for a long period of time (Onifade et al., 2007), causing alterations in soil physicochemical and microbiological properties (Ijah and Antai, 2003; Odokuma and Dickson, 2003). The overall effects of crude oil on agricultural land may be due to nutritional imbalances created by the spilled oil (Ijah et al., 2008; Chorom et al., 2010), causing reduced agricultural yield and consequently adversely affecting the socioeconomic lives of the people residing in the affected area due to high unemployment and poverty rates The toxicity of petroleum products varies widely, depending on the source which determines the composition, concentration and environmental factors. Petroleum hydrocarbons can cause extensive damage to the soil environment and to the organisms living in that particular ecosystem. The components of the hydrocarbon have been reported to be carcinogenic, mutagenic, potent immunotoxicants and neurotoxic. These components are able to accumulate in living tissues thus posing a serious threat such as death or mutation in off springs (Ilyina *et al.,* 2003).

The serious damage done by petroleum oil to the environment had led to a concerted effort of isolating and characterizing bacteria that can survive in that environment. Bacteria can only grow on nutrients or substrate that it has affinity for and can metabolically break and process. Petroleum hydrocarbon is mainly composed of carbon and hydrogen. Bacteria exposed to hydrocarbons become adapted by acquiring the ability (enzymes) to utilize petroleum hydrocarbon as a sole source of energy and carbon for the purpose of metabolic activities. Those bacteria break petroleum’s components, derive nutrients from it and render the contaminants innocuous (Ranjan *et al.,* 2014; Islam *et al.,* 2013).

Petroleum oil is the major source of energy worldwide and the balance of the ecosystem relies on each organism (micro and macro organism); thus the ability of certain life to exist on this substance is of outmost importance. In this study, we attempt to identify bacteria which can utilize crude oil as carbon and energy source from crude oil contaminated soil.

### PROBLEM STATEMENT

Petroleum oil is the most prominent source of energy for vehicles, trains, plane, and ships. It is also used for household purposes. As the use of petroleum expands with industrialization, petroleum oil becomes a greater potential source of contaminants in the soil and water environments (Bento *et al*., 2003). Contamination of soils, groundwater, sediment, surface water and air with petroleum is a major problem facing the world today. Certain diseases namely cancer, bone marrow damage and kidney and liver diseases have been associated with exposure to high concentration of petroleum oil (Olukunle, 2013).

The effects of oil pollution can be fatal and devastating not only to human, but also to animal and agricultural lands. Some animals, such as birds, mammals, and fish can be killed if petroleum hydrocarbons are ingested. Many may die from eating oil contaminated prey. Birds and insects may die if oil coats the feathers and wings, as that will prevent free locomotion and the ability to stay warm. Furthermore, oil also causes the water to have a bad odor and leave a sticky film on the surface of water making it unsafe and unpleasant to consume (Farrington and McDowell, 2004; Jain *et al*., 2011). It is therefore necessary to find solutions to solve these environmental problems.

### JUSTIFICATION OF STUDY

Crude oil exploration, exploitation and handling have been associated with frequent oil spill resulting from oil pipeline vandalization/rupture, equipment failure, tanker accidents and indiscriminate disposal of products. Tanee and Kinako (2008) have reported that despite more stringent environmental regulations, the risk of an oil spill affecting ecosystems, devastating biodiversity and stripping soils of nutrients is still high and must be accepted as inevitable; necessitating the choice of this project which is less destructive and less expensive. Therefore, this project has also been selected so as to know the organisms that can be used in bioremediation and to degrade hydrocarbonic pollutants in the environment.

### OBJECTIVES OF THE STUDY

To determine the degradative potentials of bacterial species isolated from crude petroleum oil contaminated soil.

#### Specific Objectives

- To isolate as many as possible culturable crude oil degrading bacteria from crude oil contaminated soil.
- To identify the isolated bacterial strains to species level using morphological features and biochemical characteristics
- To determine the biodegradation potential of all isolated bacterial strains singly under controlled laboratory conditions to select the best crude oil degraders.

### SCOPE OF THE STUDY

Samples were collected from three communities (Emeyal 1, Imiringi and Elebele) contaminated with crude petroleum oil spills due to corrosion and bursting of underground pipes, vandalization of pipelines in Ogbia Local Government Area of Bayelsa State, Nigeria.

## MATERIALS AND METHODS

### COLLECTION OF SAMPLING

Three sub-surface crude oil contaminated soil samples will be collected from three communities (Emeyal II, Elebele, Emeyal I) making it a total of nine samples in Ogbia metropolis. The samples will be put in clean containers and labelled according to their respective communities, all samples will be returned to the laboratory immediately for analysis.

### CULTURE MEDIA AND COMPOSITION

The following culture media were used:

- Nutrient agar
- Mineral salt medium (MSM) liquid composed per liter (pH 7.2): NH_4_NO_3_, 4.0g; Na_2_HPO_4_, 2.0g; KH_2_PO_4_, 0.53g; K_2_SO_4_, 0.17g, MgSO_4_.7H_2_O, 0.10g, and trace element solution (per 100ml); EDTA 0.1g, ZnSO_4_ 0.042g, MnS04 0.178g, H_3_BO_3_ 0.05g, NiCl 0.1g, solid MSM was prepared by adding agar (20 g/L).
- All media were sterilized by autoclaving at 121 °C for 15 minutes.

### PREPARATION OF MEDIA

- Nutrient agar was prepared following the manufacturer’s guide. 28g of nutrient agar powder was weighed into a conical flask, 1000ml of distilled water is added. Media was sterilized at 121°C for 15 minutes.
- Mineral salt agar was prepared by weighing the aforementioned salts (with some modifications) into a conical flask containing agar and dissolving in 1000ml of distilled water. Media was autoclaved at 121°C for 15 minutes.

### ISOLATION OF BACTERIA

One gram of soil sample was suspended in 9ml sterile distilled water and was diluted serially up to 10^-5^. 1ml aliquot of the suspension was inoculated separately in to nutrient agar plate using pour plate method. The plates were incubated at 37°C for 24hours, plate counting was done afterwards and recorded. After 24hours of incubation, the discrete bacteria colonies on each of the plates were counted, using a digital colony counter, model: S-961 and the plates having colonies within the range of 30 - 300 were calculated for their colony forming unit (CFU) per gram (g). The CFU/g was calculated thus:

*cfu*/*g* = *M* × *N* ÷ *D*M: Number of colony count; N: amount of sample plated on the petri dish (1ml); D: dilution factor. After 24hour incubation, predominant bacteria colonies on nutrient agar were sub-cultured using nutrient agar plates to get pure culture.

### SREENING OF THE ISOLATES BASED ON GROWTH AND HYDROGEN DEGRADATION

Pure isolates from the previous step were inoculated on mineral salt medium by spread plate method, sterile peptone water was used as diluent. Indeed, the best crude oil degrading isolates were purified on solid MSM agar plates containing crude oil, as the sole carbon and energy source, by spread-plate technique. The plates were wrapped with aluminum foil, incubated in the dark at 30 °C for 2 weeks. Colonies forming clear zones on the coated solid MSM medium were selected. After 2 weeks of incubation, individual isolates on the MSM-crude oil enriched media were subcultured on nutrient agar for 24 hours for bacterial identification.

### IDENTIFICATION AND CHARACTERIZATION OF BACTERIAL ISOLATES

Bacterial isolates were characterized using different cultural, microscopic, and biochemical characteristics. Identification of bacterial isolates was achieved by comparing their characteristics with those of known taxa

#### Gram staining

Gram staining is a method that differentiates bacteria in two large group namely gram positive and gram negative. This method differentiates bacteria by the chemical and physical properties of the cell walls. Peptidoglycan which is a polymer composed of polysaccharide and peptide chains is thick in gram positive and thin in gram negative bacteria. A Gram positive test results in a purple/blue color while a Gram negative results in a pink/red color depending on whether crystal violet or methylene blue was used. To prepare bacteria for gram staining, a smear of bacteria taken from a pure culture was made. The smear was made by mixing bacteria colony with distilled water on a dry and clean glass slide using a sterile loop. The smear was allowed to air dry, and then passed two to three times through the flame for fixation without exposing the slide directly to the flame. The smear was treated with methylene blue solution for one minute and it was washed using distilled water. The slide was then flooded with gram’s iodine kept for one minute and washed. The slide was flooded with gram’s decolorizer (95% alcohol), kept for 30 second and washed properly. A counter stain (safranin) was added for 60 second and rinsed with distilled water. The slide was blotted dry with the laboratory’s tissue paper. The slide was then observed under the light microscope at 100x objective with oil immersion (Tortora, Funke and Case, 2014).

#### Biochemical test

Biochemical tests comprising oxidase, catalase, citrate, indole, motility, coagulase, starch hydrolysis and carbohydrate fermentation test.

##### • Indole test

This test detects the ability of cells to split tryptophan to indole using the enzyme tryptophanase. This reaction is detected by the addition of Kovac’s reagent, which contains 4 (P)-dimethylamino-benzaldehyde. This reacts with the indole to produce a red colored compound. Tryptone broth was prepared according to manufacturer’s instructions and autoclave sterilized. The broth (3ml) was dispensed into well labelled sterile universal bottles and allowed to cool to room temperature (29°C). A colony of the pure culture was inoculated into the bottle and incubated at 37°C for 24hours. After 24hours of incubation, 0.5mls of Kovac’s reagent was added to the bottle. The bottle was gently rocked. The formation of red surface layer over the yellow tryptone broth was detected as a positive result while no red surface layer was reported as a negative result (Cheesbrough, 2006).

##### • Oxidase test

Oxidase test was used to determine the presence of bacterial cytochrome oxidase enzyme using the oxidation of substrate “tetramethyl-P-phenylenediaminedihydrochloride” to indophenol a dark purple colored end product. Oxidase test was used to differentiate the families of Vibrionaceae and Pseudomonadaceae (oxidase positive) from the Enterobacteriaceae (oxidase negative). It was also used for the identification of *Acinetobacter*spp. And *Klebsiella* spp. From nutrient agar. Strip of oxidase detection was prepared on a Petri dish by placing a drop of oxidase reagent on filter paper. Then, a colony of pure culture was transferred to the strip using sterile applicator stick. The strip was observed in five seconds for the formation of deep blue or violet color as a positive result. No change of color was detected as a negative result (Cheesbrough, 2006).

##### • Citrate test

The citrate test identifies the use of citrate as a sole carbon source in the absence of other nutrients in the medium. The end products cause the bromo-thymol blue indicator in the medium to turn from forest green to royal blue. The Simmons citrate agar was prepared in the Universal bottles according to manufacturer’s instruction. The slant agar was prepared and allowed to cool at room temperature (29°C) after autoclaving. A colony of pure culture was streaked on the citrate agar slant. Then, the agar was incubated for 18 to 24 hours at 37°C. After incubation, the changes of green color of Simmon’s citrate agar to blue color was indicated as positive result and no changes of color (maintained in green) was detected as negative result (Cheesbrough, 2006).

##### • Catalase test

Catalase is an enzyme that splits hydrogen peroxide into water and oxygen. The principle of this test is to detect the presence of catalase in the microorganisms. It was used in the presumptive identification of Staphylococcus spp. A colony of pure culture was transferred to a clean and dry glass slide, well labelled, using an inoculating loop. Then, a drop of 3% hydrogen peroxide, H2O2, was placed on the colony. The production of bubbles was detected as positive result, within a few seconds. No bubbles were detected as negative result within the same time frame (Cheesbrough, 2006).

##### • Coagulase test

This was used to determine the ability of cells to produce coagulase, which causes plasma to clot by converting fibrinogen to fibrin. This test was used of the identification of *Staphylococcus aureus*. A colony of pure culture of the organism was emulsified on a loop full of distilled water placed on a clean glass slide. The slide was well labelled. Two loop-full of plasma was added to the suspension and mixed gently. The formation of clumps after 10seconds was detected as a positive result. The absence of clumping after 10seconds was detected as a negative result. All negative results were confirmed using the tube test. The tube test was performed by adding 0.8ml of the test broth in a sterile glass tube and pipetting 0.2ml of plasma into the tube. The broth was mixed gently and incubated at 37°C. The broth was examined for clothing after one hour. If no clothing was observed, it is further examined after three hours and finally after twenty (24) hours. The absence of clothing of periodic examination was detected as confirmation for negative for coagulase test (Cheesbrough, 2006).

##### • Motility test

A semi-solid agar medium was prepared in a tube, organisms were inoculated by single stab down the center of the tube to about half the depth of the medium and incubated under conditions favouring motility (37°C), tubes were examined at interval after 6hours and 1 and 2 days. Non-motile bacteria generally give growths that are confined to the stab line, have sharply defined margins and leave the surrounding medium clearly transparent, while motile bacteria typically give diffuse, hazy growth that spread throughout the medium rendering it slightly opaque. *Escherichia coli*, gram negative Enterobacteriaceae, *Enterococcus casseliflavus* and *Enterococcus gallinarum*.

## RESULT AND DISCUSSION

My sample sites (Elebele, Emeyal I and Emeyal II) will be defined by the following key

Elebele = site A

Emeyal I = site B

Emeyal II = site C

### BACTERIAL ENUMERATION

After 24hours of incubation, the discrete bacteria colonies on each of the plates were counted, using a digital colony counter to get the colony forming unit per gram (cfu/g). Table 2 shows the total viable count which is the concentration of microorganisms (bacteria) in the various sites, the values obtained were similar to those obtained by Ducklow *et al*., (1979), Bukola *et al* (2011), Nwiyi and Amaechi, (2011) who enumerated the total viable count of the organisms after plating first on nutrient agar. The lowest bacterial concentration occurred at Emeyal II sample 1 (1.23x10^6^) and Emeyal I sample 1 (6.0x 10^6^) for dilutions 10^4^ and 10^5^ respectively while the highest concentration occurred at Emeyal I sample 2 (2.23x 10^6^) and Emeyal I sample 2 (2.01x 10^7^) for dilutions 10^4^ and 10^5^respectively.Table 3 shows the mean and the standard error of the means of total viable count

**Table 1:** total viable count of microorganisms in the soil (cfu/g)

| Sample site | Sample code | Total viable count |  |
| --- | --- | --- | --- |
| | | $10^6$ | $10^7$ |
| Elebele (A) | A1 | 1.32 | 1.01 |
|  | A2 | 2.15 | 1.68 |
|  | A3 | 1.45 | 1.06 |
| Emeyal I(B) | B1 | 1.75 | 0.60 |
|  | B2 | 2.23 | 2.01 |
|  | B3 | 2.04 | 1.98 |
| Emeyal II(C) | C1 | 1.23 | 1.00 |
|  | C2 | 1.84 | 1.38 |
|  | C3 | 1.49 | 1.40 |

**Table 2:** shows the mean and standard error of mean.

| Sample site | No of Sample | Total viable count |  |
| --- | --- | --- | --- |
| | | $10^6$ | $10^7$ |
| Elebele (A) | 3 | $1.64 \pm 0.26$ | $1.25 \pm 0.22$ |
| Emeyal I (B) | 3 | $2.01 \pm 0.14$ | $1.53 \pm 0.47$ |
| Emeyal II (C) | 3 | $1.52 \pm 0.18$ | $1.26 \pm 0.13$ |

All values are means ± standard errors.

**Table 3:** shows the sample sites and the number of isolates isolated from each site.

All values are means ± standard errors. **Table 3:** shows the sample sites and the number of isolates isolated from each site.
| Sample site | Sample code | Isolates |
| --- | --- | --- |
| Elebele (A) | A1 | A1<sub>1</sub>, A1<sub>2</sub>, A1<sub>3</sub>, A1<sub>4</sub> |
|  | A2 | A2<sub>1</sub>, A2<sub>2</sub>, A2<sub>3</sub>, A2<sub>4</sub> |
|  | A3 | A3<sub>1</sub>, A3<sub>2</sub>, A3<sub>3</sub>, A3<sub>4</sub> |
| Emeyal 1 (B) | B1 | B1<sub>1</sub>, B1<sub>2</sub>, B1<sub>3</sub>, B1<sub>4</sub>, B1<sub>5</sub> |
|  | B2 | B2<sub>1</sub>, B2<sub>2</sub>, B2<sub>3</sub> |
|  | B3 | B3<sub>1</sub>, B3<sub>2</sub>, B3<sub>3</sub>, B3<sub>4</sub> |
| Emeyal 2 (C) | C1 | C1<sub>1</sub>, C1<sub>2</sub>, C1<sub>3</sub>, C1<sub>4</sub>, C1<sub>5</sub>, C1<sub>6</sub> |
|  | C2 | C2<sub>1</sub>, C2<sub>2</sub>, C2<sub>3</sub>, C2<sub>4</sub> |
|  | C3 | C3<sub>1</sub>, C3<sub>2</sub>, C3<sub>3</sub>, C3<sub>4</sub>, C3<sub>5</sub>, C3<sub>6</sub> |

### DEGRADATIVE STRENGTH OF ISOLATES ON MSM-CRUDE OIL MEDIUM

Colonies forming clear zones due to the utilization of the crude oil were selected as hydrocarbon degraders. Fortunately, all 40 isolates were hydrocarbon degraders as they were isolated from crude oil contaminated soil, but some organisms were stronger degraders compared to others and these organisms are similar to those isolated by Tebyanian *et al*., 2013 and their strength of degradation was seen as follows:

**Figure 1:**
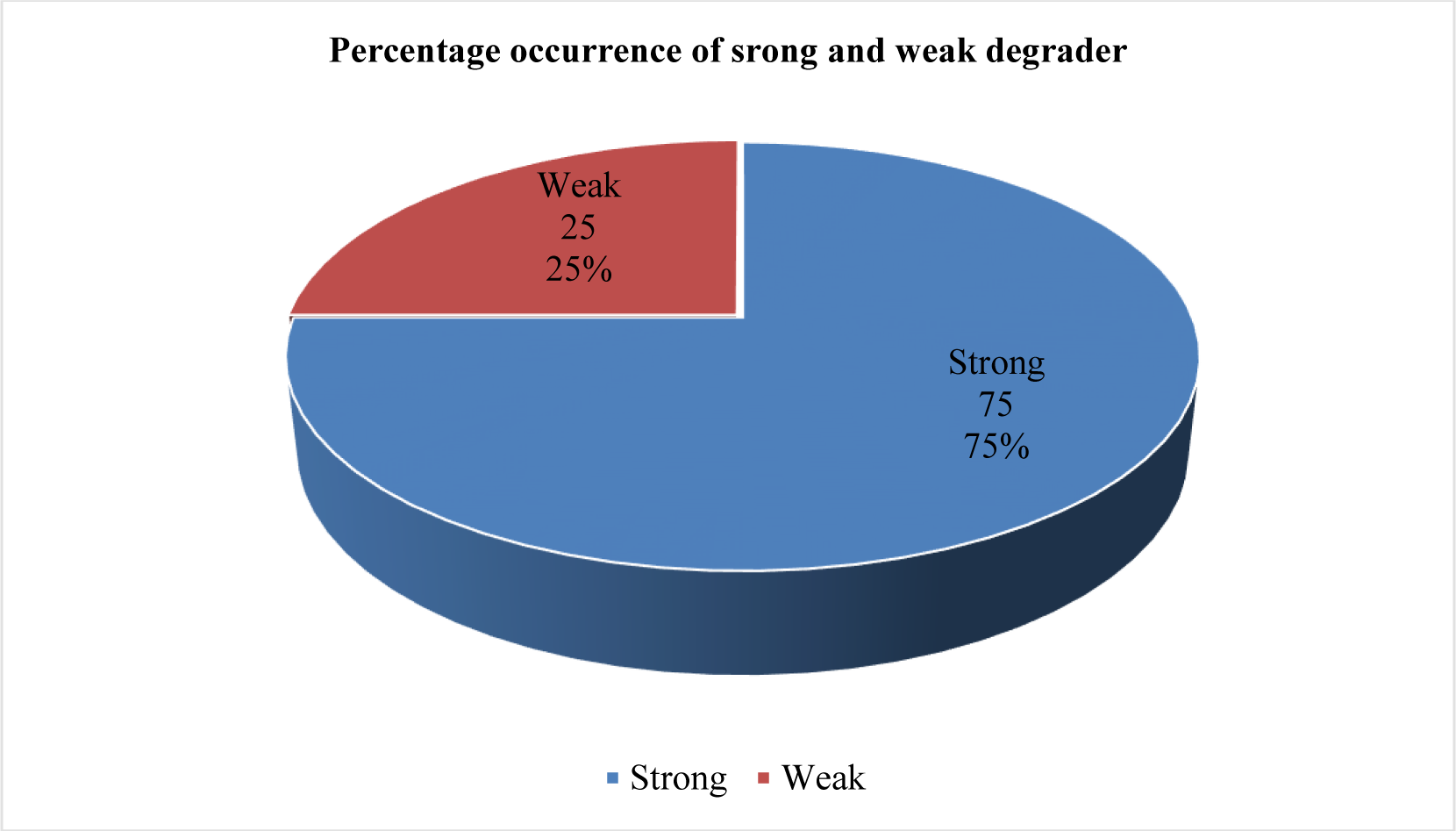
A pie chart showing the percentage occurrence of strong and weak degraders.

After 14 days’ incubation of inoculated MSM-crude oil enriched medium, the isolates were further subcultured on unenriched nutrient agar at 37°C for 24 hours for bacterial identification (gram staining and biochemical test) as seen in table 5. The bacteria isolated included *Pseudomonas* spp, *Bacillus* spp, *Escherichia coli*, *Arthrobacter* spp, *Enterobacter* spp, *Acinetobacter* spp, CON *Staphylococcus* spp, *Staphylococcus aureus*, *Klebsiella* spp, *vibrio* spp, *Micrococcus* spp.

**Table 4:** shows the degrading strength of the individual isolates.

|  | Isolates |  |  |  |  |  |
| --- | --- | --- | --- | --- | --- | --- |
| Strength | A1<sub>1</sub> | A1<sub>2</sub> | A1<sub>3</sub> | A1<sub>4</sub> |  |  |
|  | ++ | ++ | ++ | ++ |  |  |
| Strength | A2<sub>1</sub> | A2<sub>2</sub> | A2<sub>3</sub> | A2<sub>4</sub> |  |  |
|  | ++ | ++ | + | + |  |  |
| Strength | A3<sub>1</sub> | A3<sub>2</sub> | A3<sub>3</sub> | A3<sub>4</sub> |  |  |
|  | ++ | ++ | ++ | ++ |  |  |
| Strength | B1<sub>1</sub> | B1<sub>2</sub> | B1<sub>3</sub> | B1<sub>4</sub> | B1<sub>5</sub> |  |
|  | ++ | ++ | ++ | ++ | + |  |
| Strength | B2<sub>1</sub> | B2<sub>2</sub> | B2<sub>3</sub> |  |  |  |
|  | + | ++ | ++ |  |  |  |
| Strength | B3<sub>1</sub> | B3<sub>2</sub> | B3<sub>3</sub> | B3<sub>4</sub> |  |  |
|  | ++ | ++ | ++ | ++ |  |  |
| Strength | C1<sub>1</sub> | C1<sub>2</sub> | C1<sub>3</sub> | C1<sub>4</sub> | C1<sub>5</sub> | C1<sub>6</sub> |
|  | ++ | ++ | ++ | + | + | ++ |
| Strength | C2<sub>1</sub> | C2<sub>2</sub> | C2<sub>3</sub> | C2<sub>4</sub> |  |  |
|  | + | + | ++ | + |  |  |
| Strength | C3<sub>1</sub> | C3<sub>2</sub> | C3<sub>3</sub> | C3<sub>4</sub> | C3<sub>5</sub> | C3<sub>6</sub> |
|  | ++ | ++ | ++ | ++ | ++ | + |

KEY: + WEAK
++ STRONG

**Table 5:**
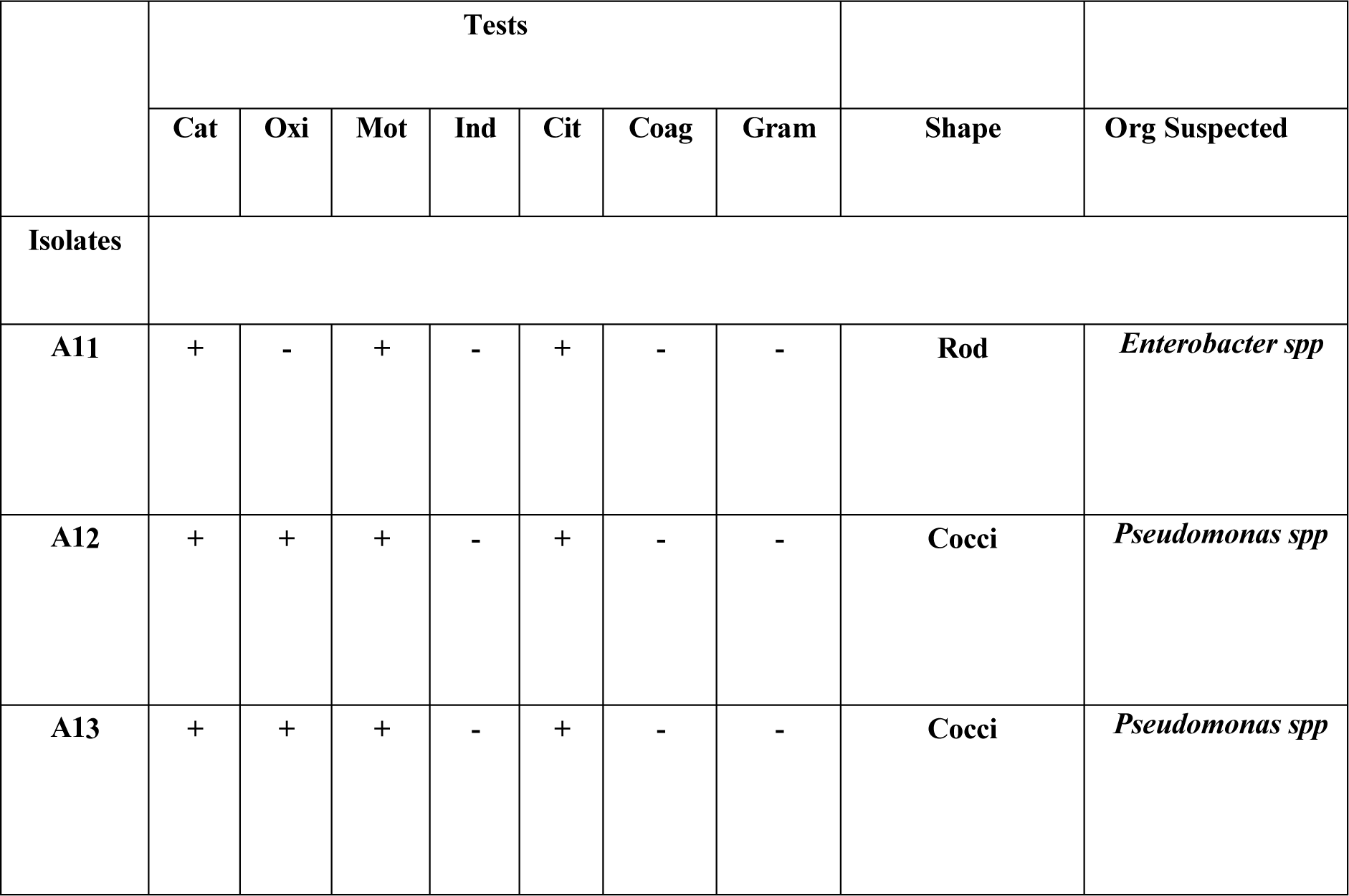
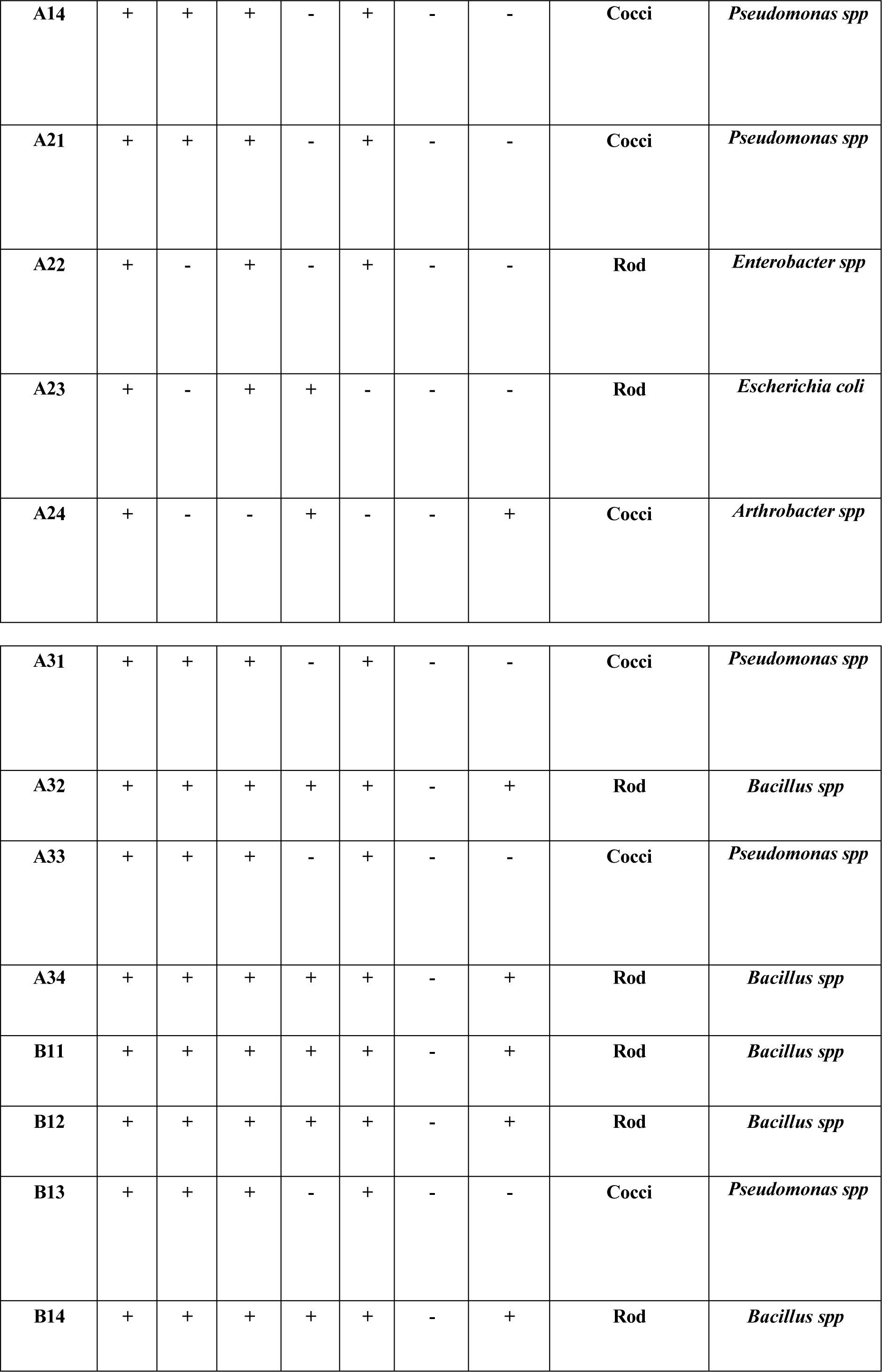
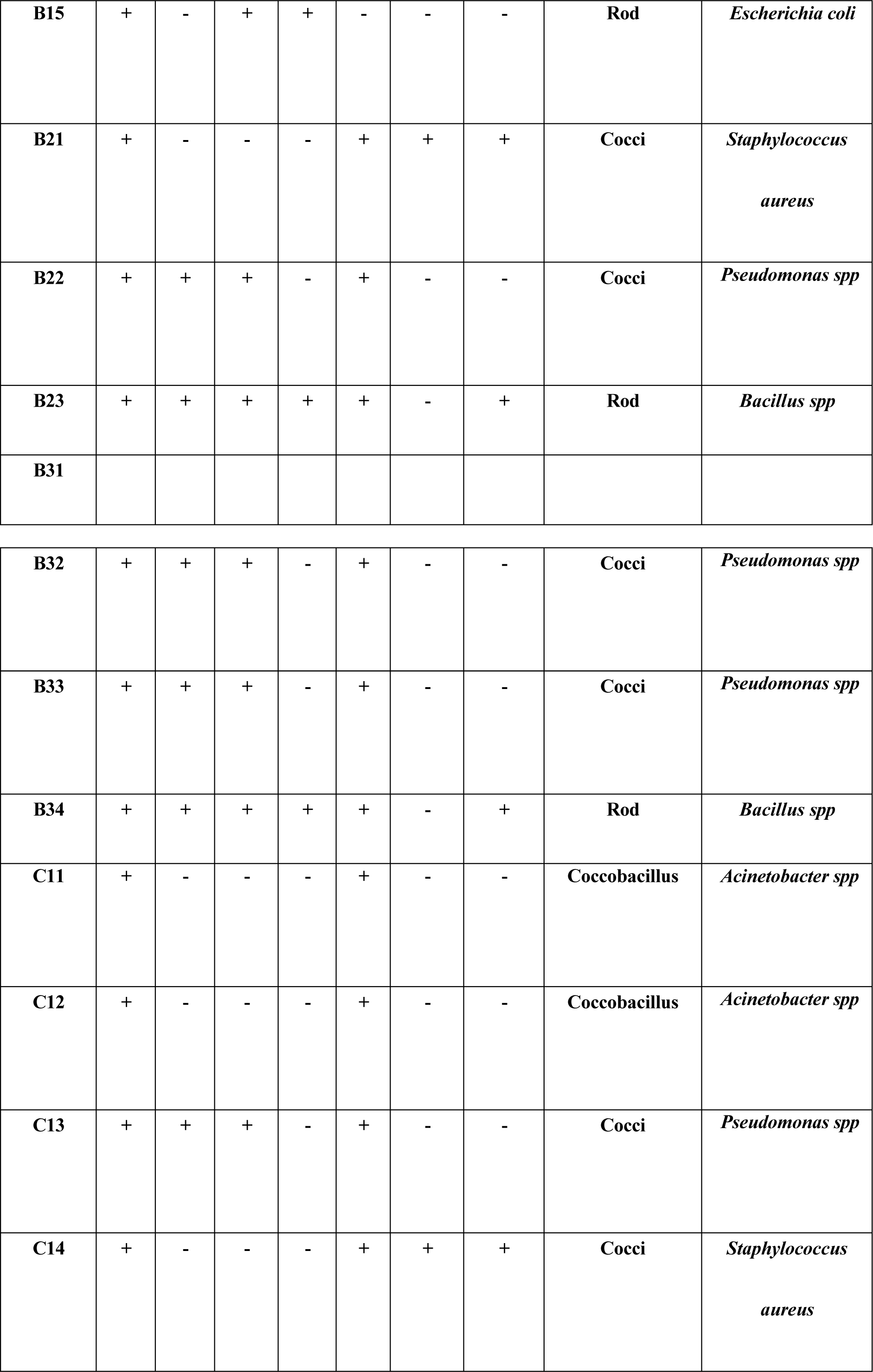
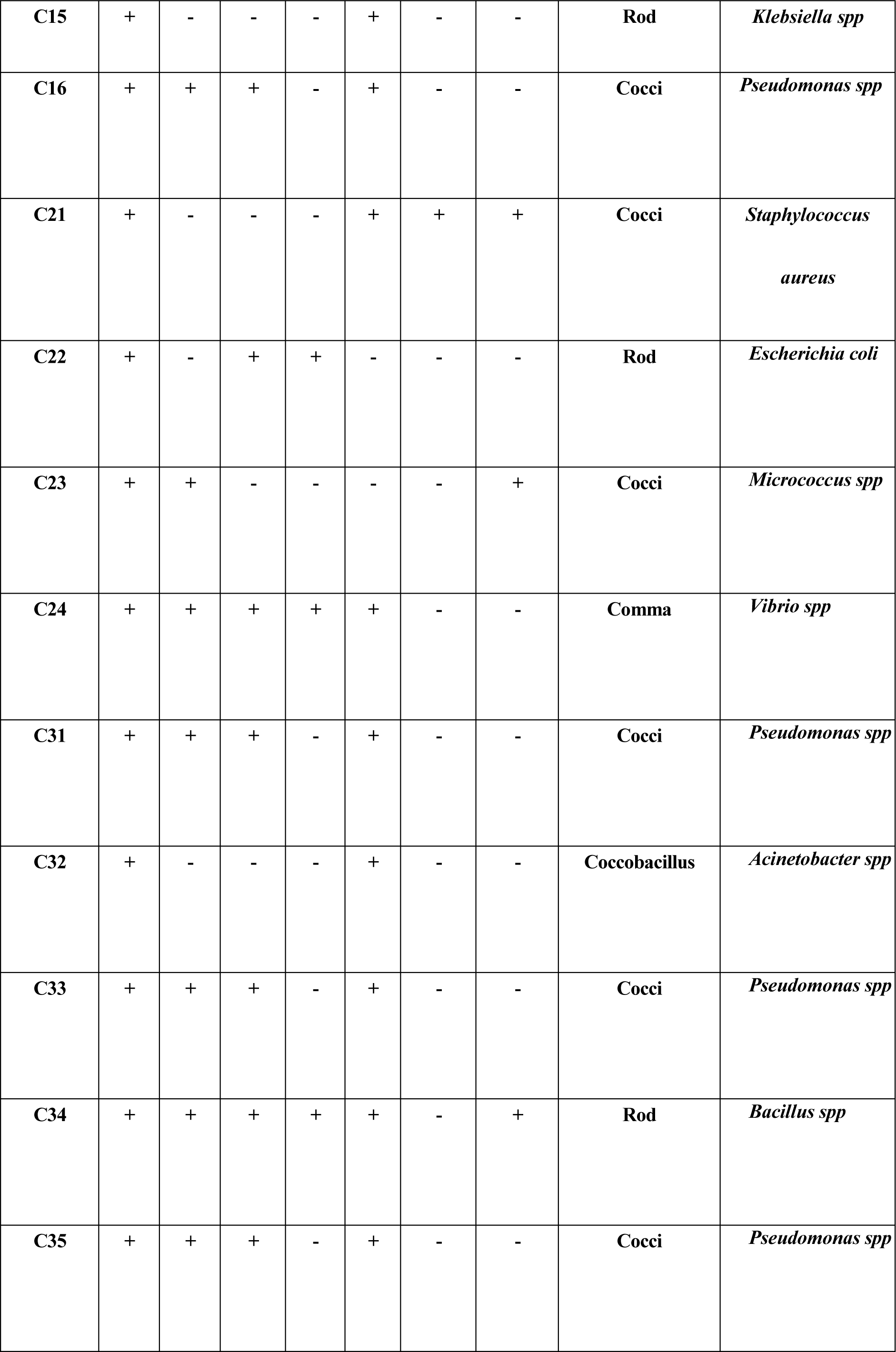
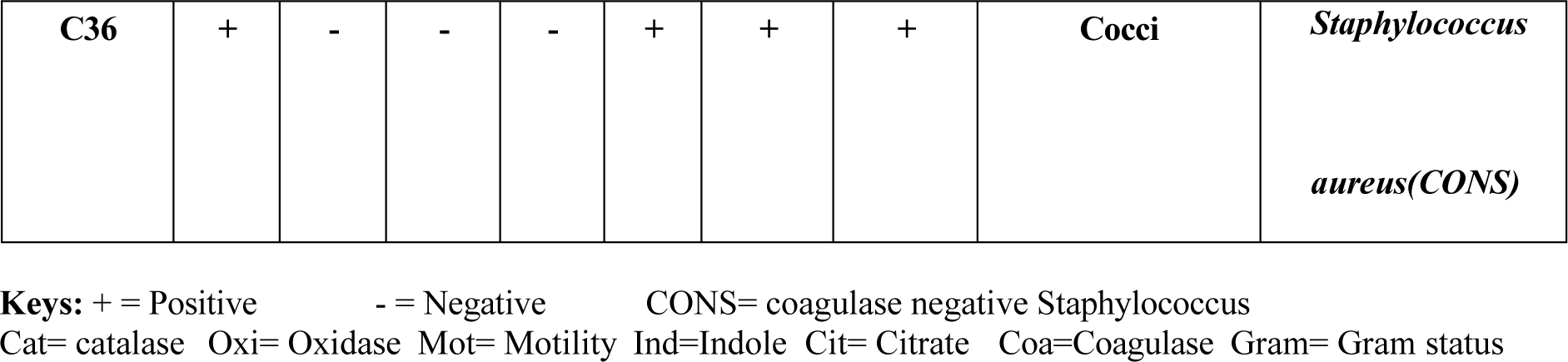
shows the gram and biochemical tests.

Ability of *Pseudomonas* to efficiently take up alkanes has been linked to production of rhamnolipid biosurfactants as was described in a syudy carried out by Sharma and coworkers (2015) using *Pseudomonas aeroginosa* DSVP20.the metabolic versatility of *Pseudomonas* has been linked to the presence of degradative plasmids such as OCT (octane), ALK (alkanes), TOL (toluene), XYL (xylene) and NAH (naphthalene) (Silva *et al*., 2006). The efficiency of *P. aeroginosa* in hydrocarbon degradation has been attributed to passive diffusion of the hydrocarbon across the cell membrane.

One specie that demonstrated growth when cultured in crude oil belonged to the genus *Klebsiella. Klebsiella* spp are well known in degradation of petroleum compounds. Among 45 hydrocarbon degrading isolates obtained from estuary sediments, Rodrigues and co-workers (2009) reported that bacteria of the genus *Klebsiella* were the most frequent encountered making 46.7% with some of them recording about 90% degradation of toluene, xylene, nonane and naphthalene.

Studies on alkane oxidation by members of the genus *Acinetobacter* have indicated that alkane utilization is widespread among this group (Ratajczak et al., 1998). In a study conducted by Mahjoubi and co-workers (2013), Acinetobacter spp was found to be the most abundant group. Similarly, Chaineau and co-worker (1999) reported that *A. baumaanii* was able to greatly assimilate saturated and aromatic hydrocarbons. Efficiency of *Acinetobacter* spp in utilization of hydrocarbon could be attributed to their ability to produce biosurfactant as was observed in a study conducted by Barkay and co-workers (1999).

### OCCURRENCE OF BACTERIAL ISOLATES

The bacteria isolated from the crude oil contaminated soil are majorly gram negative organisms as shown by the figure below. They include *Pseudomonas* spp, *Escherichia coli*, *Acinetobacter* spp, *Enterobacter* spp, *Salmonella* spp, *Vibrio* spp and *Klebsiella* spp. There was 37.5% occurrence of *Pseudomonas* spp, 7.5% occurrence of *Enterobacter* spp, *Escherichia coli* and *Acinetobacter*spp, 2.5% occurrence of *Arthrobacter*spp, *Klebsiella* spp, *Micrococcus* spp, *Vibrio* spp. *Pseudomonas* spp was dominant in all sites suggesting it is a very strong hydrocarbon degrader. The gram negative organisms found are similar to those found by Cipinyte *et al*., 2011, Erdogan *et al.,* 2011, Luo *et al.,* 2012, Nwaogu *et al.,* 2008, Zhanga *et al.,* 2018, also reported are *Bacillus* spp, coagulase negative *Staphylococcus*, *Staphylococcusn aureus.* The result reiterates the importance of these organisms to bioremediation of oil spills.

**Figure 2:**
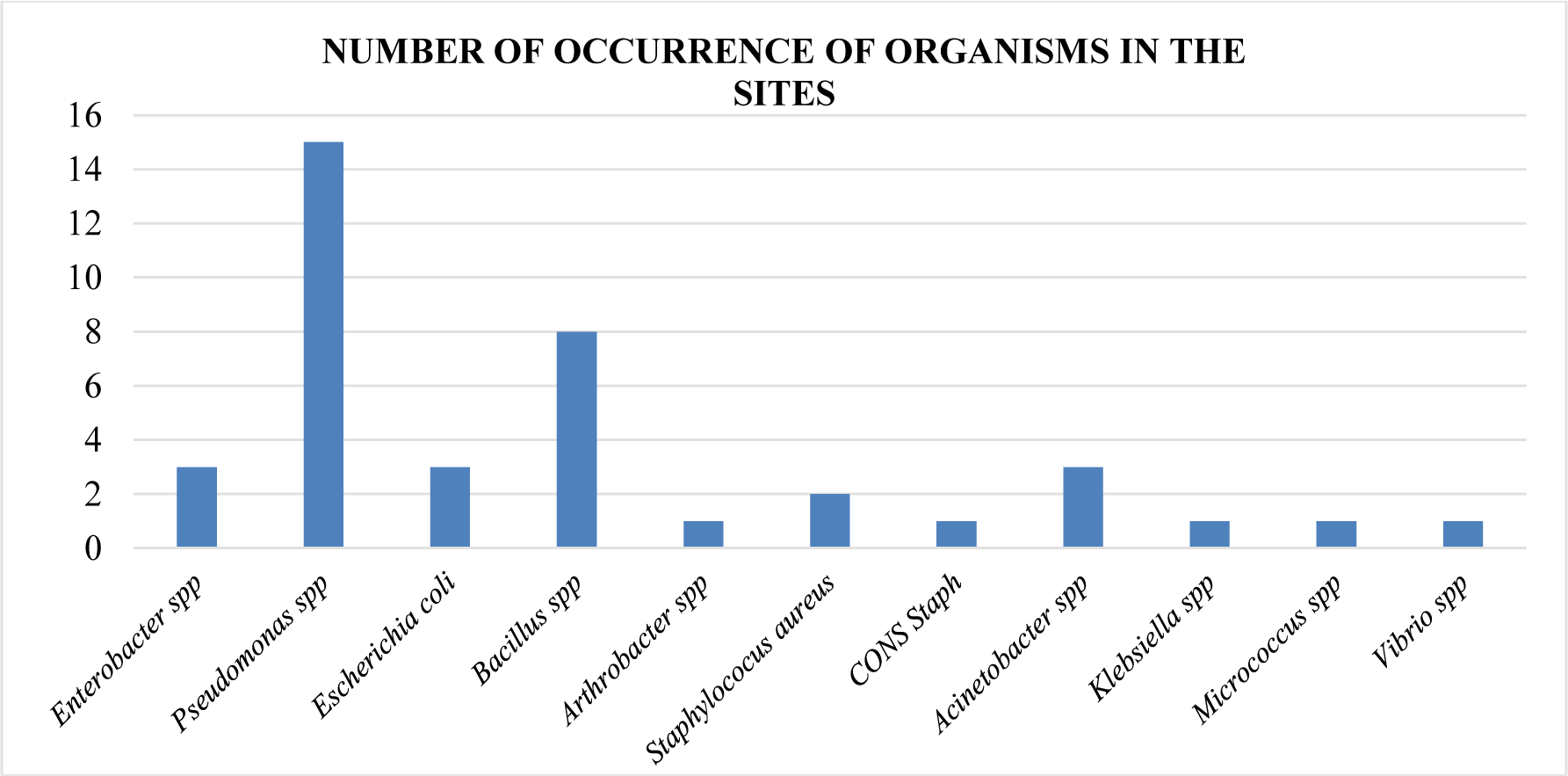
A histogram occurrence of bacteria isolates from the contaminated sites.

**Table 6:** pooled mean count (10^4^cfu/g) of bacteria from the sample sites. Elebele Emeyal I, Emeyal II.

| Elebele | Emeyal I | Emeyal II |
| --- | --- | --- |
| 7.1±1.21 | 8.7±2.39 | 7.1±0.72 |

All values are mean± standard error

## CONCLUSION

Hydrocarbon compounds, as products of the oil industry and related activities, are considered serious environmental pollutants. Bioremediation is one of the most rapidly growing areas of environmental biotechnology, by microorganisms that have specific metabolic capacities for cleaning up pollutants. The application of bioremediation is distinguished by its low costs and simple implementation with safer environmental impacts and public acceptability. It is evident from this study that hydrocarbon degrading organisms are abundant in Ogbia local government and they are isolated from hydrocarbon polluted sites. A screening program was designed to isolate interesting hydrocarbon degrading strains and in this case a combination of growth parameters and biological activities resulted the isolation of forty bacterial strainsin Elebele, Emeyel I, Emeyal II in Ogbia local government, from these, thirty efficient degraders were isolated and identified based on morphological and biochemical tests. These organisms were observed to use crude oil as their sole source of carbon. The result of this study showed that bacteria can grow in crude oil contaminated soils as ordinarily would have been thought otherwise and bacteria with this special ability are called hydrocarbonoclastic bacteria and these organisms have potentials to be used as remediation agents. The study also took into consideration the various locations where the soils were collected. The result also showed that Emeyal I contains higher and longer pollution of the environment as aging also increases the number of organisms on the contaminated soil. The high number of *Pseudomonas* spp found in the sites makes it a strong degrader in crude oil contaminated soils. The result also revealed that there are more gram negative hydrocarbon degraders than are gram positives, gram negative hydrocarbon degraders includes organisms from the genus *Pseudomonas, Flavobacterium, Escherichia, Acinetobacter, Enterobacter, Salmonella, Vibrio* etc and gram positives includes *Bacillus, Nocardia, Staphylococcus, Arthrobacter, Corynebacteria, Micrococcus* etc.

## APPENDIX

### Appendix 1: Materials used

- **Beaker:** A flat bottom vessel with a lip, used as a laboratory container mainly for the purpose of stirring, mixing, and the collections of samples.
- **Graduated Measuring Cylinder:** This is a glass ware piece of laboratory equipment with a narrow cylindrical shape it is used for critical measuring of liquid volumes in a routine experiment.
- **Wire loop/Inoculation loop**: Used to inoculate (introduce) test samples into culture media for bacterial and fungal cultures. It is reheated by flaming to red hot before use
- **Personal protective equipment’s**: These are protective clothing, designed to protect the body from injury, infection and other bio-hazardous materials e.g. hand gloves, nose guard, and eye glass. Etc.
- **Bunsen Burner**: It is laboratory equipment that produces a single open gas flame, which is used for heating sterilization and combustion
- **Electronic weighing balance**: An electronic device for measuring weight.
- **Autoclave**: A device in which pressure is used to produce high temperature steam to achieve sterilization. Autoclaving at 121°c for 15 minutes is required for sterilization. Or it is a device used for sterilization of glass wares and media.
- **Refrigerator**: This is an electronic appliance that is used to cool and preserve sample or specimen to minimize the risk of bacterial contamination and explosion of volatile materials.
- **Petri-dishes/Agar plate**: It is used to act as a supporting container to hold the culture medium for isolation and cultivation of specimen microbes.
- **Incubator**: A device used for the incubation of cultured microorganisms. It is mostly used for bacterial and fungal culture.
- **Lighter/Matches and Gas (Flame)**: A device used to ignite fire.
- **Trowel**: A gardener’s tool, shaped like a scoop, used in taking off plants, soil particles, and stirring soil etc.
- **Distilled water**: Water that has many of its impurities removed through distillation.
- **Ethyl-alcohol:** Colorless chemical liquid compound with a slight characteristic odor used as an antibacterial disinfectant for sterilization of animate and non-animate objects.
- **Cotton wool:** A soft mass of cotton, used especially for applying liquids (ethyl alcohol) for cleaning and sterilization of laboratory bench and other objects.
- **Colony counters**: A device used for counting colonies of bacteria growing in a culture.
- **Nutrient Agar**: A general purpose medium used for the cultivation of a wide variety of microorganisms, supporting the growth of non-fastidious organisms and also use for microbial count.
- **Mineral salt agar(MSM):** this media is used for isolation of hydrocarbon utilizing bacteria.
- **Test tubes**: Test tubes are widely used in the laboratory to hold, mix, or heat small quantities of solid or liquid chemicals. They are used for serial dilution in the microbiology laboratory.
- **Pipette**: A small sterile glass tube use to transport a measured volume of liquid, often as a media dispenser.
- **Microscope slide (Glass slide):** This is a thin flat piece of glass used to hold objects for examination under a microscope.
- **Soil:** It is the material found on the surface of the earth that is composed of organic and inorganic materials.
- **Polythene bag:** A plastic film bag used for containing, packaging and transporting of materials.
- **Immersion oil:** It is used to increase the resolving power of a microscope.
- **Microscope:** An instrument used to view microscopic objects (i.e. objects that cannot be seen with the naked eye).
- **Soil auger:** A hand held or rotary powered tool with a helical cutting edge used for drilling holes in soil. Augers are used for taking soil samples, drilling for caissons.
- **Spatula:** A small hand-held flat, flexible stainless steel utensils, used for mixing, spreading, lifting and transferring of materials e.g. Agar medium.
- **Cover slip:** A cover slip is a thin flat piece of transparent material that is placed over objects for viewing with a microscope.
- **Electronic soil pH meter:** This an electronic device used for measuring the acidity or basicity of the soil.
- **Aluminum foil:** This is aluminum metal prepared in thin metal leaves with a thickness less than 0.2mm. it is used for items packaging and act as a total barrier to light, oxygen, germs and moisture.
- **Mastering tape:** This used for labelling of sample for proper identification.
- **Greased pencil:** This is a wax writing tool also known as wax pencil it is used for marking on hard, glossy non-porous surface.

### Appendix 2

**a. Preparation of Nutrient agar**

Nutrient agar is a general purpose medium used for the cultivation of microbes supporting the growth of a wide range of non-fastidious organisms.

Composition:

0.5% peptone

0.3% beef/yeast extract

1.5% agar

0.5% NaCl

Distilled water

pH adjusted to neutral (7.4) at 25°C

#### Procedure

- Suspend 28g of nutrient agar powder in 1 litre of distilled water.
- Heat this mixture while stirring to fully dissolve all components.
- Autoclave the dissolved mixture at 121°C for 15 minutes.
- Once the nutrient agar has been autoclaved allow it to cool but not solidify.
- Dispense nutrient agar into plates and leave plates on the sterile surface until the agar has solidified.
- Store plates in refrigerator.

**b. Preparation of mineral salt medium**

#### Composition

Mineral salt medium (MSM) liquid composed per liter (pH 7.2): NH4NO3, 4.0g; Na2HPO4, 2.0g; KH2PO4, 0.53g; K2SO4, 0.17g, MgSO4.7H2O, 0.10g, and trace element solution (per 100ml); EDTA 0.1g, ZnSO4 0.042g, MnS04 0.178g, H3BO3 0.05g, NiCl 0.1g, solid MSM was prepared by adding agar (20 g/L).

#### Procedure

- Weigh appropriate grams of each salt and dissolve in 1 litre of de-ionized water.
- Autoclave at 121°C for 15 minutes.
- Allow media to cool but not solidified.
- Dispense the media into plates and allow to solidify.
- Store plates in refrigerator.

### Appendix 3

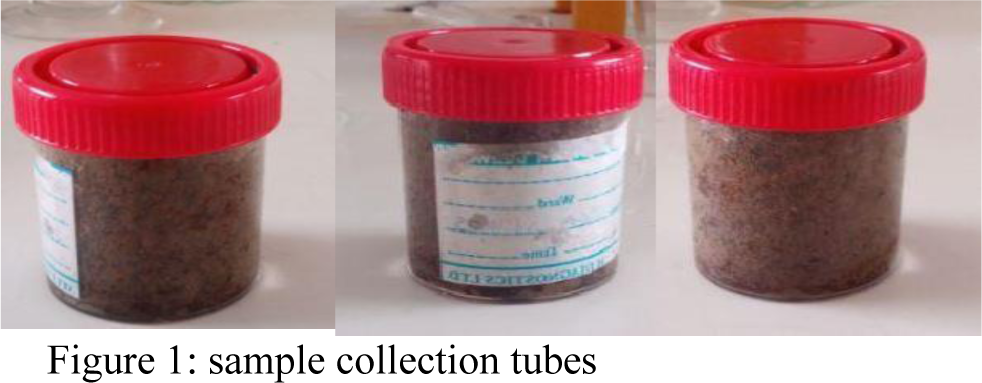

